# Comparing basal dendrite branches in human and mouse hippocampal CA1 pyramidal neurons with Bayesian networks

**DOI:** 10.1101/2020.03.14.991828

**Authors:** Bojan Mihaljević, Pedro Larrañaga, Ruth Benavides-Piccione, Javier DeFelipe, Concha Bielza

## Abstract

Pyramidal neurons are the most common cell type in the cerebral cortex. Understanding how they differ between species is a key challenge in neuroscience. A recent study provided a unique set of human and mouse pyramidal neurons of the CA1 region of the hippocampus, and used it to compare the morphology of apical and basal dendritic branches of the two species. The study found inter-species differences in the magnitude of the morphometrics and similarities regarding their variation with respect to morphological determinants such as branch type and branch order. We use the same data set to perform additional comparisons of basal dendrites. In order to isolate the heterogeneity due to intrinsic differences between species from the heterogeneity due to differences in morphological determinants, we fit multivariate models over the morphometrics and the determinants. In particular, we use conditional linear Gaussian Bayesian networks, which provide a concise graphical representation of the independencies and correlations among the variables. We also extend the previous study by considering additional morphometrics and by formally testing test whether a morphometric increases or decreases with the distance from the soma. This study introduces a multivariate methodology for inter-species comparison of morphology.

## 1 Introduction

Pyramidal neurons are the most common neuronal type in the cerebral cortex. A key challenge in neuroscience is to understand how these neurons differ across species^1–5^. They are often compared in terms of their dendritic morphology, a feature which directly influences neuronal computation^6–8^. However, dendritic morphology varies across layers and cortical regions within a single species^4,9–13^ (more so in primates than in rodents^1,2^), possibly as an adaptation to their different cytoarchitectonic and functional features. It is thus important to compare neurons from homologous regions of the two species. For the human and the mouse, one such region is the hippocampus, considered to be one of the most evolutionary conserved archicortical regions^14^.

A recent study^15^ compared morphologies of pyramidal cells’ dendrites from the hippocampus CA1 area of human and mouse brains, using the first data set of digitally reconstructed morphologies of human CA1 pyramidal cells. Because the dendritic trees in the sample were incompletely reconstructed, the study focused on branch-level morphometrics, namely on branch length, surface, volume and (average) diameter. The authors found differences in the magnitude of the morphometrics, with the dendritic branches of human neurons notably larger and thicker than those of the mouse. Human terminal branches were also non-proportionally longer than non-terminal branches, showing that the human neurons were not scaled versions of mouse neurons. Nonetheless, the authors found similarities regarding how the morphometrics varied with respect to morphological determinants such as branch type (whether a branch is terminal or not). For example, terminal branches were longer than non-terminal ones in both species.

Comparing morphometrics between species is not straightforward, because the morphometrics can vary, within a species, with respect to determinants such as branch type and branch order. In addition, some differences in determinants may be artifacts of the reconstruction process; for instance, removing incompletely reconstructed branches could yield different proportions of terminal and non-terminal branches in the two samples. For example, if our human data set had a much higher proportion of non-terminal branches than the mouse one, a test might not reject the null hypothesis of equal length, while separate tests for terminal or non-terminal branches would have (because non-terminal branches are shorter than terminal ones in both species; human non-terminal branches are longer than mouse non-terminal ones; and human terminal branches are also longer than mouse terminal branches^15^). Besides missing existing differences, the converse is also possible and inter-species that were detected with a single test might be insignificant once we account for the determinants. The authors of Ref.^15^ accounted for the determinants precisely by splitting the branches into terminal and non-terminal ones, and further down according to branch order, and then testing hypotheses of location difference (Kruskal-Wallis test) between pairs of obtained subsets of branches (e.g., mouse terminal branches of branch order two are as long as human terminal branches of the same branch order).

Instead of splitting the branches according to the determinants and running multiple tests, we could specify a multivariate statistical model over the morphometrics and the morphological determinants. Models such as Bayesian networks (BNs^)16-18^ can represent the probabilistic relationships among the variables of a domain, while algorithms for learning Bayesian networks from data can uncover such relationships. In the above example we would have three random variables —species, branch type and branch length— and learning a Bayesian network from the data would tell us that the species variable and the length variable are marginally independent yet dependent given branch type. Probabilistic dependence does not imply different locations (e.g., mean or median), and a morphometric with the same mean yet different variance in the two species is indeed dependent on the species variable. Thus, only after identifying a dependence between the species variable and a morphometric (with the Bayesian network) we would proceed to test for location difference between species. In addition, Bayesian networks can simplify comparison as they provide a concise graphical representation of the branch-level morphology, in terms of probabilistic relationships among morphometrics and determinants. This representation allows us to easily identify independencies and correlations among the morphometrics and determinants and thus identify whether a particular difference is due to intrinsic inter-species differences, due to differences in determinants, or due to a combination of both.

In this paper, we use the hippocampus cells used in Ref.^15^ (henceforth ‘the previous study’) yet focus on basal dendrites alone. In addition to branch length and average branch diameter, we consider three additional branch-level morphometrics: tortuosity, bifurcation angles and tilt angles. We study how these morphometrics vary according to four morphological determinants —species, branch type, branch order, and Euclidean distance from the soma. We learn from data Bayesian network models for the joint distribution over the morphometrics and the determinants. In particular, we use conditional linear Gaussian Bayesian networks (CLGBNs), a model family that allows us to jointly model discrete and continuous variables. The conditional regression models contained within the CLGBNs allow us to identify the magnitude and sign of correlations. We perform Kruskal-Wallis tests of location difference regarding branch tortuosity, and bifurcation and tilt angles, the morphometrics that were not covered in the previous study. An additional contribution, not present in Ref.^15^, is that we formally test whether a morphometric increases or decreases with the distance from the soma, by assessing linear correlation. While the authors of Ref.^15^ reported such observations (e.g., branch diameter decreased with increasing branch order in non-terminal branches), they did not test for correlation between variables and based such claims on the rejection of the hypothesis of equal median diameters across branch orders. As mentioned above, we focus on basal dendrites and thus leave apical dendrites for future work.

We learn Bayesian networks from three different subsets of our data set: (a) from terminal branches alone; (b) from non-terminal branches alone; and (c) from both terminal and non-terminal branches. Separating the data into (a) and (b) lets us learn models that are specific to each branch type, whereas (c) allows us to study the differences between terminal and non-terminal branches. For each data subset (that is, (a), (b), and (c)), we learn a separate Bayesian network model for each species, as well as a combined Bayesian network model for both species. The species-specific models give insight into the specifics of the species whereas the combined ones allows us identify the independencies and correlations with the species variable.

## 2 Methods

### 2.1 Data

All neuron morphology reconstructions were obtained by the authors of Ref.^15^ by intracellular injections in the CA1 region of the hippocampus. There were 54 human neurons, obtained at autopsy from two donors both no apparent neurological alterations (a male aged 45 and a female aged 53) within a postmortem interval of 2–3 hours. Upon removal, the brains were immersed in cold 4% paraformaldehyde in 0.1 M phosphate buffer, pH 7.4 (PB). There were 50 mouse neurons, obtained from nine C57BL/6 adult (8-week-old) male mice. All animals were overdosed with sodium pentobarbital and perfused though the heart with phosphate-buffered saline (0.1 M PBS) followed by 4% paraformaldehyde in PB. 3D coordinates of the dendritic morphology were extracted using Neurolucida 360 (MicroBrightfield, VT, USA; see Figure 1a). Since intracellular injections of the cells were performed in coronal sections, the part of the dendritic arbor nearest to the surface of the slice from which the cell soma was injected (typically at a depth of ~ 30*μm* from the surface) was lost. Thus, some branches were not fully included within a section and were therefore incompletely reconstructed. For further details on tissue reconstruction, intracellular injections, immunocytochemistry, and cell reconstruction, see Ref.^15^.

**Figure 1.**
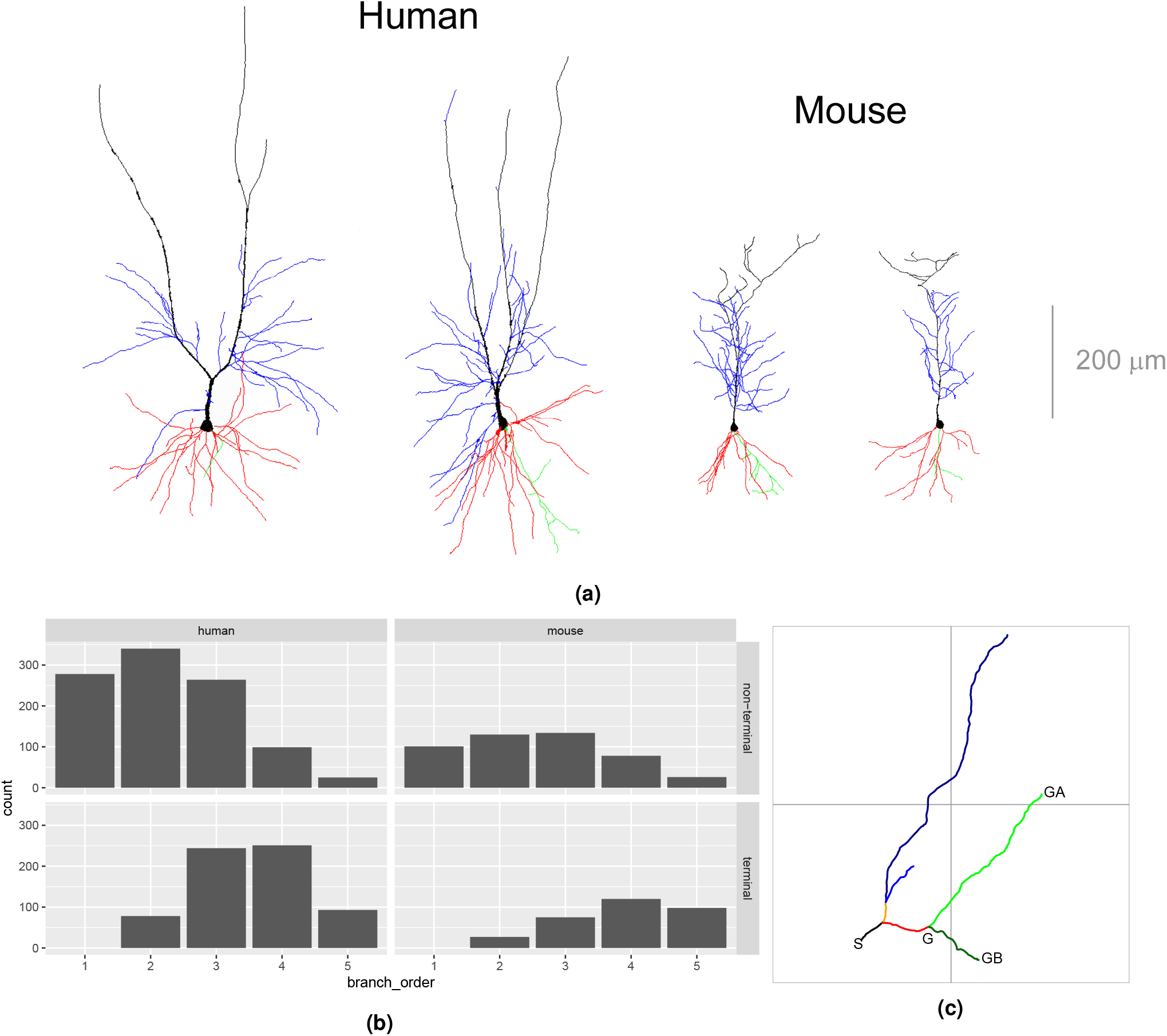
(a) Two human (left) and mouse (right) pyramidal cells’. Basal arbor is shown in red; main apical dendrite in black; apical collateral dendrites in blue, and axons in green (adapted from Ref.^15^). (b) Branch frequencies per species, branch type, and branching order. (c) An illustration of the computed morphometrics and morphological determinants. A branch is a sequence of straight lines (i.e., segments) between consecutive bifurcation points or between a bifurcation point and a terminal point (each branch shown with a different color). Branch orders (variable `branch_order`) are computed centrifugally, with branch order 1 corresponding to branches emanating from the soma (the black branch). The length (`length`) of a branch is the sum of the lengths of its segments. The average diameter (`diameter`) of a branch is the weighted average of the diameters of the branches’ segments. There are two types of variables (variable `branch_type`): non-terminal branches (black, red and orange branches) and terminal. The distance from the soma (`distance`) is the length of the straight line from the dendrites’ insertion point into the soma up to the starting point of a branch (e.g., for the green branches, this is the distance between S and G). Variables `distance, length`, and `diameter` are all measured in micrometers (*μm*). `tortuosity` describes dendritic meandering and is defined as the ratio of `length` and the length of the straight line between the beginning and the end of a branch; it is a unitless quantity and `tortuosity` = 1 denotes a perfectly straight branch whereas increasing values denote more tortuous branches. Remote bifurcation angle (`bifurcation_angle`) is the shortest planar angle between the vectors from the bifurcation to the endings of the daughter branches; for the red branch, it is the smaller angle between points GA, G, and GB. Remote tilt angle (`tilt_angle`) is the smallest angle between the branch director vector and the vectors from the bifurcation to the endings of the daughter branches. `bifurcation_angle` and `tilt_angle` are measured in radians. `bifurcation_angle` and `length` are undefined for terminal branches, as they do not bifurcate.

We only considered branches from branch order 1 to branch order 5, as only 4% of the branches corresponded to higher branching orders. We omitted from our analysis 23 branches that had a single child branch, as well as one trifurcating branch, because angles are only defined for branches that bifurcate (i.e., have exactly two child branches); we kept the descendants of these branches. We omitted all incomplete branches from our analysis (all incomplete branches were labeled as terminal).

Our final sample consisted of 2462 branches, 1672 from 54 human neurons and 790 branches from 50 mouse neurons (see Figure 1b). There were 136 (mean) ± 95 (standard deviation) branches per each combination of species, branch type, and branching order (not considering branch order 1 for terminal branches, as there were none of this order). There were more human than mouse branches (1672 and 789, respectively) and more non-terminal than terminal branches (1475 and 986, respectively). In both species, roughly 60% of the branches were non-terminal, due to the removal of incomplete terminal branches, as half the branches of a fully reconstructed bifurcating tree are necessarily terminal and the other half non-terminal.

### 2.2 Morphometrics

Because some branches were incompletely reconstructed, we did not use morphometrics that depend on the arbor being complete, such as total arbor length or topological variables such as partition asymmetry. Instead, we focused on five branch-level morphometrics (see Figure 1c). Namely, we computed the branch length (`length`); remote bifurcation angle (`bifurcation_angle`); remote tilt angle (`tilt_angle`); average branch diameter (`diameter`); and branch tortuosity (`tortuosity`). We considered one continuous morphological determinant, namely, the Euclidean distance from the branch starting point to the soma (distance), and three discrete ones, namely: species (`species`), (centrifugal) branch order (`branch_order`), and branch type (i.e., whether terminal or non terminal; `branch_type`). Note that bifurcation angles are only defined for non-terminal branches. While the previous study analyzed branch surface and branch volume, in addition to `length` and `diameter`, we omitted these variables as they were deterministic functions of `diameter` and `length` and thus any differences between species would be implied by differences in `diameter` and `length`. We computed all morphometrics and determinants with the open-source NeuroSTR library (https://computationalintelligencegroup.github.io/neurostr/).

### 2.3 Bayesian networks

A Bayesian network (BN)^16^ 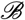 allows us to compactly encode a joint probability distribution over a vector of *n* random variables **X** by exploiting conditional independencies among triplets of sets of variables in **X** (e.g., **X** is independent of *Y* given *Z*). A BN consists of a directed acyclic graph (DAG) 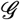 and a set of parameters 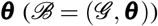. The vertices (i.e., nodes) of 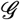 correspond to the variables in **X** while its directed edges (i.e., arcs) encode the conditional independencies among **X**. A joint probability distribution 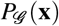 encoded by 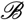, where **x** is an assignment to **X**, factorizes as a product of local conditional distributions,

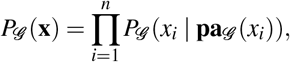

where 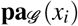 is an assignment to variables 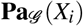, the set of parents of *X_i_* in **X** according to 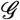 (for a continuous variable, a probability mass function 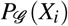 is replaced with a density function 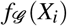). 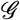 induces conditional independence constraints for 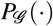, derivable from the basic constraints that each *X_i_* is independent of its non-descendents in 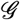 given 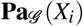. For example, for any pair of variables *X, Y* in **X** that are not connected by an arc in 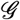 there exists a set of variables **Z** in **X** (disjoint from {*X*} and {*Y*}) such that *X* and *Y* are independent conditionally on **Z** (i.e., 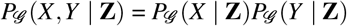). Similarly, for any pair of variables *X, Y* in **X** that are connected by an arc in 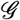 there is no set **Z** such that *X* and *Y* are independent conditionally on **Z**. These constraints extend to nodes not connected by an arc in 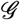 and the structure 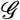 thus lets us identify conditional independence relationships among any triplet of sets of variables *X, Y*, and **Z** in **X**. For example, in the DAG *X* → *Y* → *Z* we only have one independence: *X* is independent of *Z* conditional on *Y*; *X* and *Y*, *X* and *Z*, and *Y* and *Z* are not marginally independent.

The parameters **θ** specify the local conditional distributions (densities) 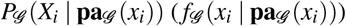 for each variable *X_i_*. When **X** contains both discrete and continuous variables, as in our case, a common approach is to let 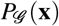 be a conditional linear Gaussian distribution (CLG)^19^. The CLG only allows discrete parents in 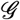 for the discrete variables **D** in **X** and therefore the local conditional distribution of each discrete variable *D_i_* in **D** is a categorical distribution. The local conditional density 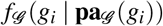 for a continuous variable *G_i_*, in the set of continuous variables **G**, is

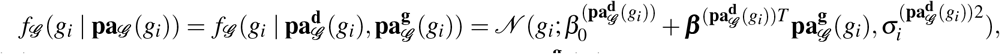

where 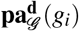 is an assignment to the discrete parents of *G_i_* and 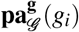 an assignment to the continuous parents of *G_i_*. There is thus a different vector of coefficients (***β***, *σ*^2^) for each assignment 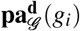 and indeed the conditional density *f*(**g** | **d**) is a multivariate normal. 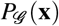 is a mixture of multivariate normal distributions over **G**, with one component for each instantiation **d** of **D**.

### 2.4 Learning Bayesian networks from data

Learning a Bayesian network 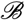 from a data set 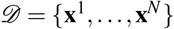 of *N* observations of **X** involves two steps: (a) learning the DAG 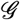; and (b) learning ***θ***, the parameters of the local conditional distributions. There are two main approaches to learning 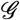 from 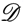^16^: (a) by testing for conditional independence among triplets of sets of variables (the *constraint-based* approach); and (b) by searching the space of DAGs in order to optimize a score such as penalized likelihood (the *score-based* approach). While seemingly very different, conditional independence tests and network scores are related statistical criteri^a20^. For example, when considering whether to include the arc *Y* → *X* into a graph 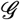, the likelihood-ratio test of conditional independence of *X* and *Y* given 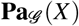 and the Bayesian information criterion^21^ (BIC) score are both functions of log 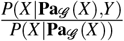. They differ in computing the threshold for determining independence: the former relies on the distribution of the statistic under the null model (i.e., conditional independence) whereas the latter is based on an approximation to the Bayes factor between the null and alternative models. Besides using different criteria, the constraint-based and score-based approaches also differ in model search, that is, in terms of the sets *X*, *Y*, and **Z** that they choose to test conditional independence for. The score-based approaches tend to be more robust^16^, as they may reconsider previous steps in the search by removing or reversing previously added arcs. A typical score-based search algorithm is hill climbing, a local search which starts from some initial DAG 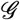 and greedily adds, removes or reverses arcs as long as that improves the score of the DAG. While classical hypothesis testing of conditional independence might be more in line with previous work on our data set^15^, score-based learning allows for more robust algorithms while still using sound statistical criteria for independence testing. We thus used the latter approach in this paper.

### 2.5 Location and goodness-of-fit tests

We complement the previous study^15^ with location tests for `tortuosity`, `bifurcation_angle`, and `tilt_angle`, morphometrics that were not considered in that paper. Although t-tests and the ANOVA fit naturally with conditional linear Gaussian Bayesian networks, we follow the previous study and apply the Kruskal-Wallis (KW) rank sum tests. We test our assumptions that the morphometrics are normally distributed conditional on `species`, `branch_type` and `branch_order` (see Section 2.6) with the Kolmogorov-Smirnov (KS) test. We set *α* = 0.05 as the significance level for all tests.

### 2.6 Assumptions

By modelling a joint distribution *P*(**x**) with a 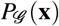 encoded with a CLGBN, we are assuming the following about *P*(**x**): (a) all conditional independencies in 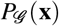 are present in *P*(**x**); (b) *f*(**g** | **d**) is a multivariate normal density for each **d**; and (c) variables in **D** have no parents in **G** in the DAG of *P*(**x**). In particular, (a) means that some dependencies implied by 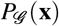 need not hold in *P*(**x**). (b) states that for each combination of `species`, `branch_type`, and `branch_order`, the distribution over `diameter, length, tortuosity, distance, bifurcation_angle`, and `tilt_angle` is a multivariate normal. (b) in turn implies that: (i) the morphometrics marginally follow a Gaussian distribution; and (ii) the dependencies among them are linear. (i) is a common simplifying assumption while we consider (ii) to a reasonable assumption given that there are not many cases (136 (mean) ± 95 (standard deviation) branches per combination of type, species, and branching order, see Section 2.1). In addition, violations of (b) do necessarily lead to wrong arcs in 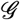, and we are precisely interested in the learning structure of 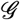 rather than the accurate estimation of *P*(**x**). Since 3 of our 9 variables are discrete, (c) means that a significant number of arcs in 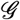 might have forced directions, as `species, branch_type`, and `branch_order` can have no parents in 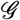.

### 2.7 Settings

As mentioned above, we consider three basic settings for learning Bayesian networks: from terminal branches alone (Section 3.2); from non-terminal branches alone (Section 3.3); and from both terminal and non-terminal branches (Section 3.4). In each setting, we learn three networks: one for the human branches; one for the mouse branches; and a combined model for branches of both species. Branching angles are only defined for non-terminal branches, and thus we omit them in models of terminal branches (Section 3.2) as well as in combined models of terminal and non-terminal branches (Section 3.4). In Section 3.4 we focus on the `branch_type` node and avoid analyzing dependencies among morphometrics, as branch-type specific dependencies can get obfuscated with the combined data from terminal and non-terminal branches that we use in Section 3.4.

All branches at branch order 1 had zero distance from the soma. To avoid a singular conditional Gaussian distribution for `distance`, we replaced these zeros by sampling from a normal distribution with mean 0 and standard deviation 0.00001.

We learned network structures by using the tabu algorithm^22^, implemented in the bnlearn^23^ R^24^ package, to optimize the BIC score. The tabu algorithm is a local search that efficiently allows for score-degrading operators by avoiding those that undo the effect of recently applied operators; we used a tabu list of size 30 and allowed for up to 30 iterations without improving network score. Since dependencies implies by the learned networks might be false positives (see assumption (a) in Section 2.6), we verify some of them with a posterior check. This consists in learning a Bayesian network by maximizing the BIC score over the variables in question only and verifying the dependence in that network.

## 3 Results

We begin by assessing whether the conditional distributions of the morphometrics can be reasonably approximated with a Gaussian distribution when conditioned on `species, branch_type` and, optionally, `branch_order` (Section 3.1). We then learn Bayesian networks from terminal branches (Section 3.2), non-terminal branches (Section 3.3), and both terminal branches and non-terminal branches (Section 3.4).

### 3.1 Morphometrics’ conditional distributions

Considering the morphometrics’ distributions conditional on species and branch type (i.e., of the form *P*(*X* | `species,branch_type`)), the KS test did not reject normality (at the 0.05 significance level) for 6 out of 20 conditional distributions (see Table 1 in the Supplementary Information). For terminal branches, this included `diameter` for both species and `length` for the mouse (see Figure 2). For non-terminals, it included `bifurcation_angle` for both species and `tilt_angle` for the mouse. When also conditioning on branch order (i.e., considering distributions of the form *P*(*X* | `species, branch_type, branch_order`)), the KS test did not reject normality for 68 out of 90 conditional distributions (see Table 2 in the Supplementary Information). This included `bifurcation_angle` and `tilt_angle` regardless of species, branch type and branch order, as well as `length` and `diameter` of terminal branches for either species. On the other hand, the KS test rejected normality at most branch orders for the `length` of non-terminal branches and `tortuosity` of human branches. In summary, the conditional distributions of most morphometrics can be approximated with Gaussian distributions after accounting for `branch_order`.

**Figure 2.**
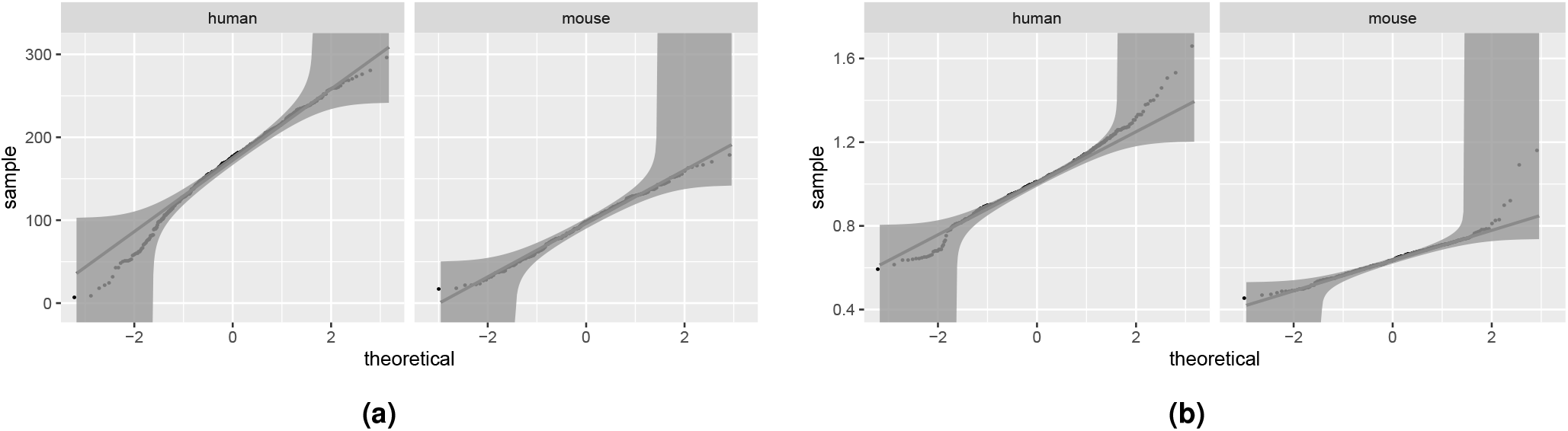
Q-Q plots of `length` (a) and `diameter` (b) of terminal branches conditional on `species` and `branch_type`. Normality is only rejected for human terminal branches, with a p-value of 0.04. The distribution of mouse `length` is especially close to the theoretical one (p-value 0.98). The plots show quantiles of the sample distributions on the Y axis and the quantiles of the standard normal distribution on the X axis. 95% KS confidence bands are shown; the hypothesis of normality is rejected at the 5% level if the sample does not fall completely within the bounds. Distributions that deviate little from normality have the points lying close to the straight line.

### 3.2 Terminal branches

The learned human and mouse BNs (Figure 3a and Figure 3b, respectively) uncovered a set of independencies among the variables. For example, `diameter` was independent of all determinants and morphometrics, including `length`, in both species, as its node was disconnected from the rest of the graph in Figure 3a and Figure 3b. `tortuosity` and `length` were marginally independent in the mouse BN and a posterior analysis found them independent in human branches as well, although the human BN did not imply this independence (due to the path `tortuosity` → `distance` → `length`).

**Figure 3.**
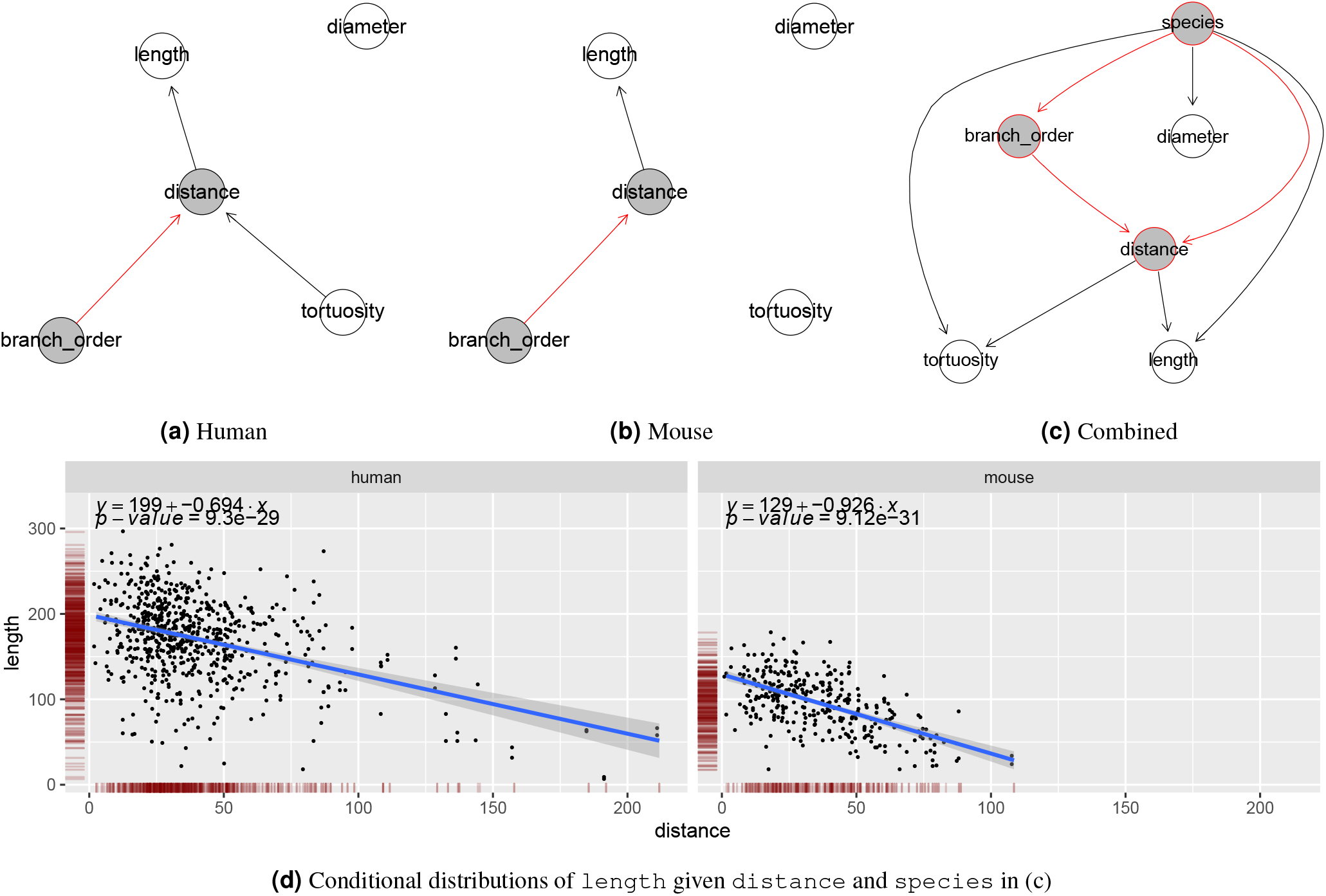
Bayesian networks learned from the terminal branches. human data (a), mouse data (b), and data of both species (c). Proximity between two nodes of a graph is unrelated to the strength of their correlation. The nodes of the morphological determinants are shaded in grey with red borders. Arcs among morphological determinants are depicted in red. (d) Scatter plots depicting the conditional distribution of `length` on `distance`, for the human (left) and for the mouse (right). The linear regression line shows the mean of the fitted conditional distribution of `length` as a function of `distance`, with its formula given in the top part of the panel, with `y` standing for `length` and *x* for `distance`. The band shows a 95% confidence interval around the mean.

The marginal correlations of the determinants (`branch_order` and `distance`) with the morphometrics were similar for the two species. Namely, `length` decreased with `distance` (see Figure 3d), with a correlation coefficient (between `length` and `distance`) *ρ* = −0.58 for the mouse and *ρ* = −0.41 for the human. In addition, length was independent of `branch_order` given distance for both species. On the other hand, as mentioned above, diameter was independent of both `branch_order` and distance. What differed between the species was that `tortuosity` slightly increased with distance in human cells (*ρ* = 0.30; note the `tortuosity` → `distance` arc) while in the mouse it stayed constant (no arcs in or out of `tortuosity`). In summary, the effect of increasing `distance` was that `length` decreased, `diameter` stayed constant, while tortuosity increased slightly in human branches yet stayed constant in the mouse ones. Naturally, distance itself increased with `branch_order` (note the `branch_order` → `distance` arc in both BNs; see also Figure 4). Overall, the human and mouse BNs were similar, with the only structural difference being that the `tortuosity` → `distance` arc was present in the human BN yet not in the mouse BN.

**Figure 4.**
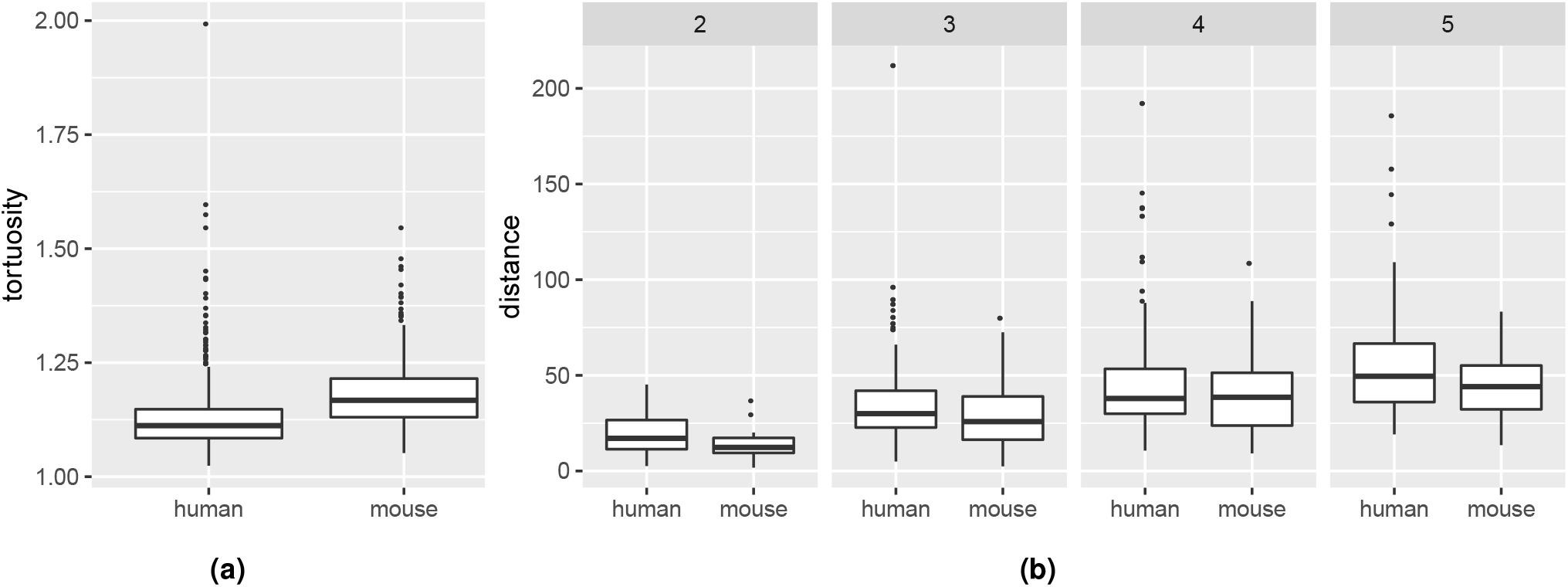
Differences in magnitude of `tortuosity` (a) and `distance` (b) of terminal branches between the species. Human terminal branches were less tortuous than mouse terminal branches (KW p-value < 0.0001 for `tortuosity`). Marginally, they were not located further away from the soma than mouse ones (p-value 0.26 for `distance`) yet, for branch orders 2, 3 and 5, they were (p-values 0.0096, 0.0091, 0.0897, and 0.0398 for `distance` at branch orders 2, 3, 4, and 5, respectively).

The combined BN for terminal branches (Figure 3c) shows that the determinants’ distributions varied between the human and mouse data sets, as there were arcs from the `species` node to both `branch_order` and `distance`. Indeed, there were proportionally more order 4 and 5 terminal branches in the mouse data set (68%) than in the human one (52%; see also Figure 1b). This was likely an artifact of incomplete reconstruction (see Figure 3 in the Supplementary Information and Section 2.1) rather than an actual difference in branching orders between the two species. On the other hand, a KW test found no difference (p-value of 0.26) in `distance` between the species. However, Figure 3c implies that `distance` does differ between `species` once we account for `branch_order` (as both species and branch_order and parents of distance in the BN) and, indeed, we confirmed that human branches were located further away from the soma than mouse branches of the same branching order (Figure 4b). This difference between species was obfuscated marginally by the higher proportion of high order branches in the mouse data set and was thus undetected by the KW test.

The combined BN for terminal branches also shows that the distributions of all morphometrics differed between species, since there was an arc from the `species` node to each morphometric. In particular, we found that human terminal branches were less tortuous than mouse terminal branches (Figure 4a); as reported by the previous study, they were also thicker and longer. The combined BN shows that these differences were not only due to differences in determinants between the two samples (e.g., proportionally more high order branches in the mouse), because the `species` node was not independent of any morphometric given any the determinants. The differences could also not be explained in terms of other morphometrics. For example, the lower `tortuosity` of human terminal branches was unrelated to their higher `length`, as `tortuosity` was independent of `length` given `distance` in the combined BN. These were, therefore, intrinsic differences between human and mouse terminal branches.

The difference in `diameter` was in fact unrelated to differences in the determinants, as `diameter` was independent of all other variables given `species`. The differences in length and `tortuosity`, on the other hand, could not be explained in terms of intrinsic inter-species differences alone, because `length` and `tortuosity` were not independent of the determinants given `species` (`distance` and `species` were parents of both `tortuosity` and `length`). Nonetheless, we found that the inter-species ratios of mean `length` and `tortuosity` were roughly constant across branch orders.

### 3.3 Non-terminal branches

The human BN (Figure 5a) and the mouse BN (Figure 5b) uncovered a number of conditional independencies among the variables. For example, in the mouse BN all morphometrics were independent: (a) of the determinants, `branch_order` and distance, given diameter; and (b) of distance given `branch_order`. In the human BN, tortuosity and `tilt_angle` were independent of all variables, including the determinants, given length and `bifurcation_angle`, respectively.

The marginal correlations of the determinants (`branch_order` and `distance`) with the morphometrics were similar for the two species. In particular, `diameter` decreased with `distance` (*ρ* = −0.35 for the human and *ρ* = −0.47 for the mouse; see Figure 1 in the Supplementary Information). While significant, the correlation of `distance` with other morphometrics was weak, with no correlation coefficient larger, in absolute value, than 0.29. Thus, `bifurcation_angle` and `tilt_angle` slightly decreased with `distance, length` slightly increased, and `tortuosity` stayed constant. The correlation coefficients were similar between the species for all morphometrics (Figure 1 in the Supplementary Information). As mentioned above, distance was independent of all morphometrics given `branch_order` in the mouse BN, while in the human BN it was only correlated with `bifurcation_angle` and `tilt_angle` given `branch_order`.

**Figure 5.**
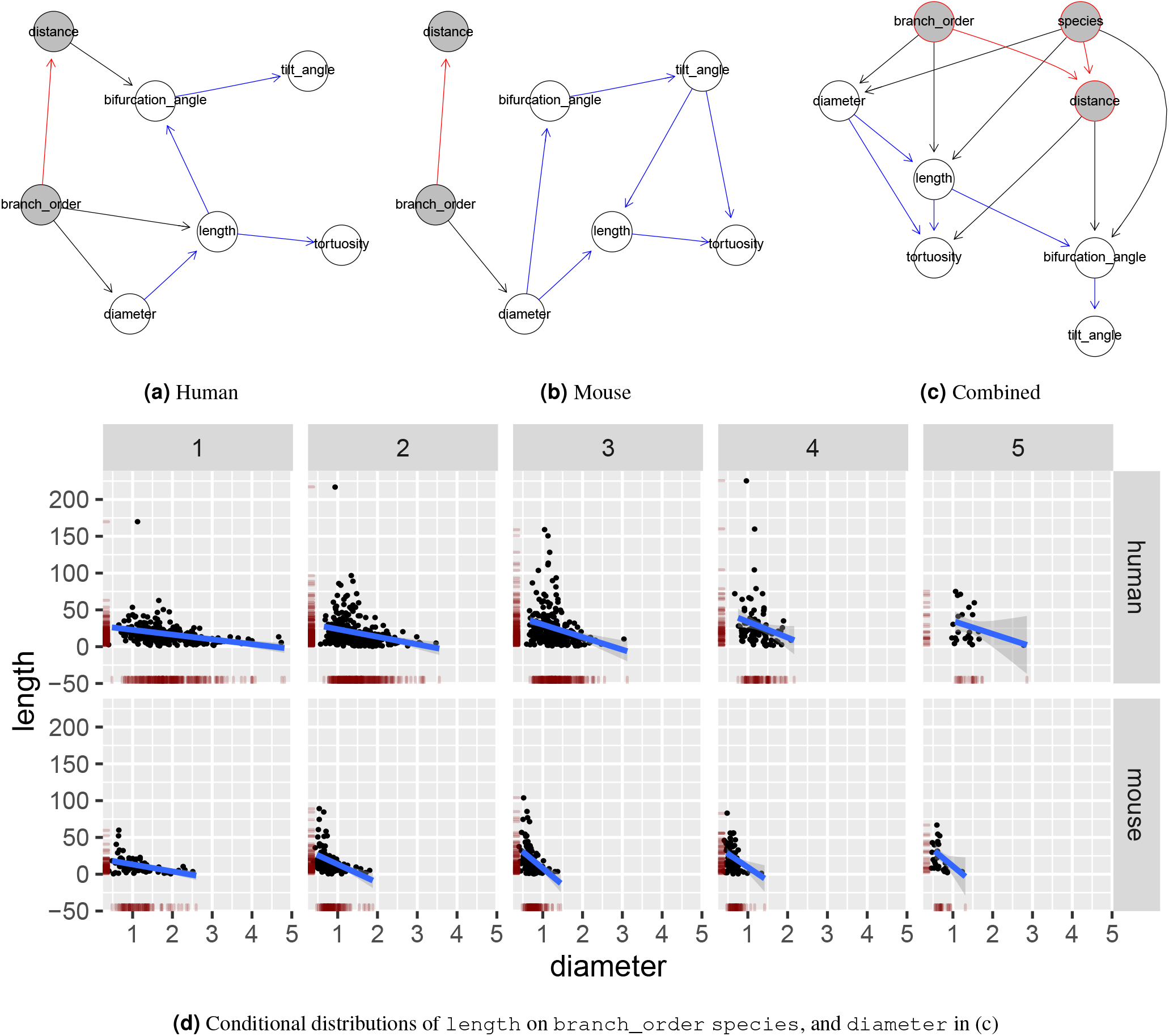
Bayesian networks learned from the non-terminal branches of human data (a), mouse data (b), and data of both species (c). Proximity between two nodes of a graph is unrelated to the strength of their correlation. The nodes of the morphological determinants are shaded in grey with red borders. Arcs among morphological determinants are depicted in red; among morphometrics in blue; the rest in black. (d) Scatter plots depicting the conditional distribution of `length` on `species, branch_order` and `diameter`. The linear regression line shows the mean of the fitted conditional distribution of `length` given `diameter`, with a 95% confidence interval; there is one regression line for each combination of `branch_order` and `species`. Since `length` is independent of `branch_order` given diameter in (b), the regression lines are roughly parallel across branch orders for the mouse.

A number of correlations among morphometrics were similar across the species. In particular, diameter and length were negatively correlated (see Figure 5d), with the correlation coefficient varying across branch orders for the human yet not for the mouse (hence no `branch_order` → length arc in the mouse BN). `tortuosity` was positively correlated with length in both species, with the longer branches being more tortuous (see Figure 2 in in the Supplementary Information). `bifurcation_angle` increased slightly with diameter (*ρ* = 0.11 for the human and *ρ* = 0.29 for the mouse) and decreased slightly with length (*ρ* = −0.17 for the human and *ρ* = −0.21 for the mouse). `tilt_angle` was positively correlated with `bifurcation_angle` in both human (*ρ* = 0.58) and mouse (*ρ* = 0.53).

Overall, the the human and mouse BNs were less similar that those for the terminal branches. There were five arcs common to the two BNs. The human BN had more arcs from the determinants to the morphometrics (black arcs in Figure 5a), while the mouse BN had more arcs among the morphometrics (blue arcs in Figure 5b).

The combined BN for non-terminal branches (Figure 5c) shows that the distribution of the `distance` determinant varied between the human and mouse data sets, while that of `branch_order` did not. As in terminal branches, a KW test found no difference (p-value of 0.2267) in `distance` between the species. However, Figure 5c implies that distance does differ between species once we account for `branch_order` and, indeed, we confirmed that human non-terminal branches were located further away from the soma than mouse non-terminal branches of the same branching order (Figure 6e). The marginal independence of species and `branch_order` in Figure 5c means that the distribution of `branch_order` did not vary significantly between the mouse and human data sets (see also Figure 1b).

**Figure 6.**
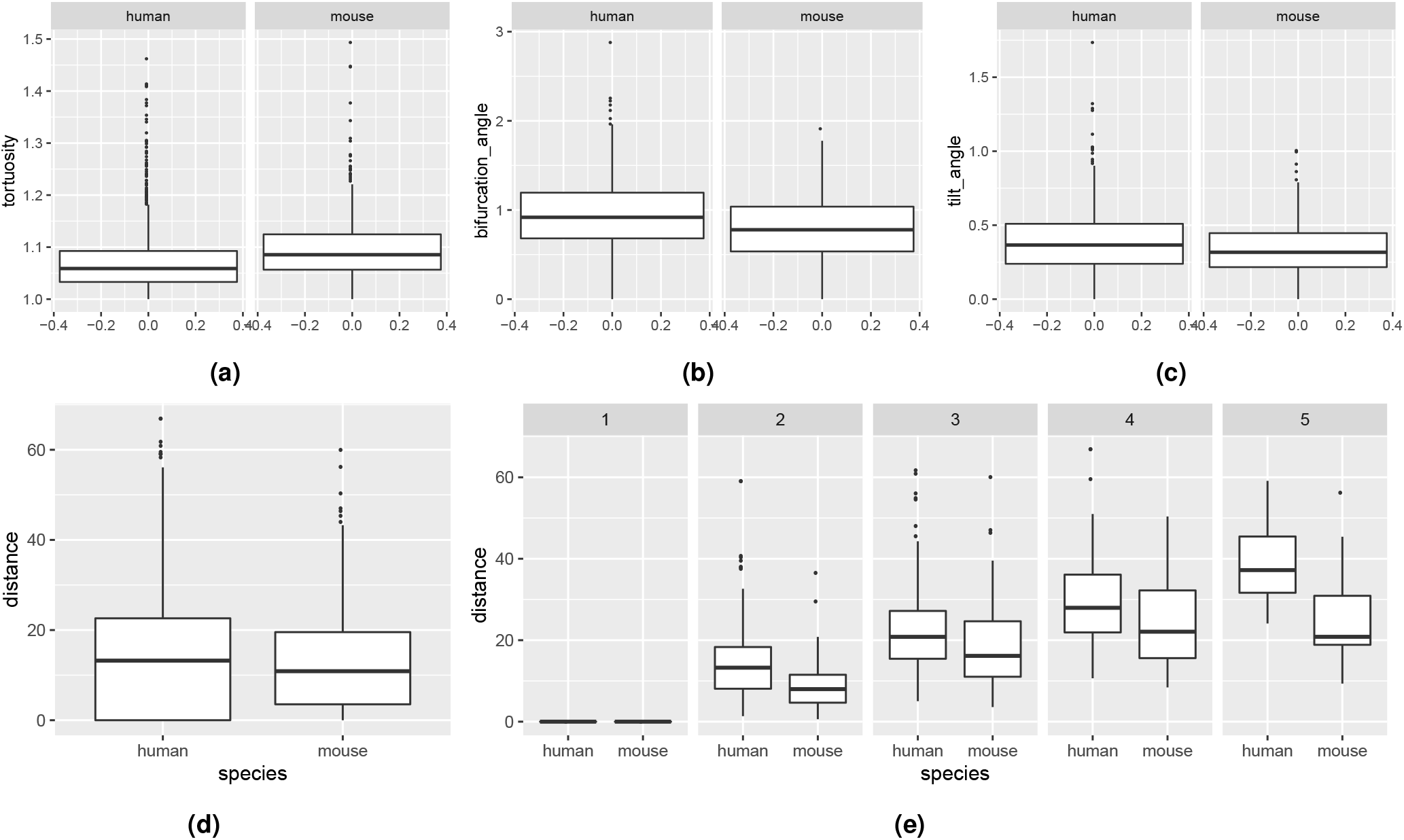
Differences in magnitude of variables of non-terminal branches between the species. Human non-terminal branches were less tortuous (a), and had larger bifurcating (b) and tilt angles (c) (KW p-values < 0.0001). Marginally, human non-terminal branches were not located further away from the soma than mouse ones ((d); p-value 0.2267 for `distance`) yet (e), for all branch orders except for 1, they were (p-value of 0.3377 at branch order 1, 0.0003 at branch order 4, and < 0.0001 at branch orders 2, 3, and 5)

The combined BN for non-terminal branches also shows that the distributions of all morphometrics differed between species. In particular, we found that human non-terminal branches were less tortuous and had larger bifurcation and tilt angles than mouse ones (Figure 6a to Figure 6c); as reported by the previous study, they were also thicker and longer than those of mouse. The differences in `tortuosity` and `tilt_angle` could be explained in terms of other morphometrics and determinants, without reference to species. Namely, `tortuosity` was independent of `species` given `length, diameter`, and distance, while `tilt_angle` was independent of species given `bifurcation_angle`.

The differences in the morphometrics could not be explained in terms of intrinsic inter-species differences alone, as no morphometric was independent of `distance` and `branch_order` given `species` (either `distance` or `branch_order` were parents for any morphometric). Indeed, the inter-species differences in `length, tilt_angle`, and `tortuosity` were less pronounced or inexistent when conditioning on branch order. In particular, differences in length were insignificant at branch orders 2, 4, and 5; in `bifurcation_angle` at branch orders 4 and 5; while the differences in `tortuosity` were only significant at branch order 2 (see Figure 4 in the Supplementary Information). This effect did not hold for `diameter` and `tortuosity` as they did differ between species after conditioning on `branch_order`.

### 3.4 All branches

The human (Figure 7a) and mouse (Figure 7b) BNs for all branches show that all variables varied with respect to `branch_type`, including the determinants, `branch_order` and distance. In particular, terminal branches were, in both species, less tortuous than non-terminal ones (KW p-values < 0.0001; see Figure 7c); as reported by the previous study, they were also thinner and longer than non-terminal branches. The differences with respect to `length, diameter`, and `tortuosity` between terminal and non-terminal branches could not be explained only in terms of the remaining determinants nor the remaining morphometrics, as `branch_type` was directly connected to all other variables in Figure 7a and Figure 7b. For example, the lower tortuosity of non-terminal branches could not be explained only by their shorter length, as tortuosity is not independent of `branch_type` given `length` in Figure 7a nor in Figure 7b. These seem to be, instead, intrinsic differences between terminal and non-terminal branches. The human and mouse BNs were similar, with the only structural difference being that the `distance` → `tortuosity` arc was present in the human BN yet not in the mouse BN.

**Figure 7.**
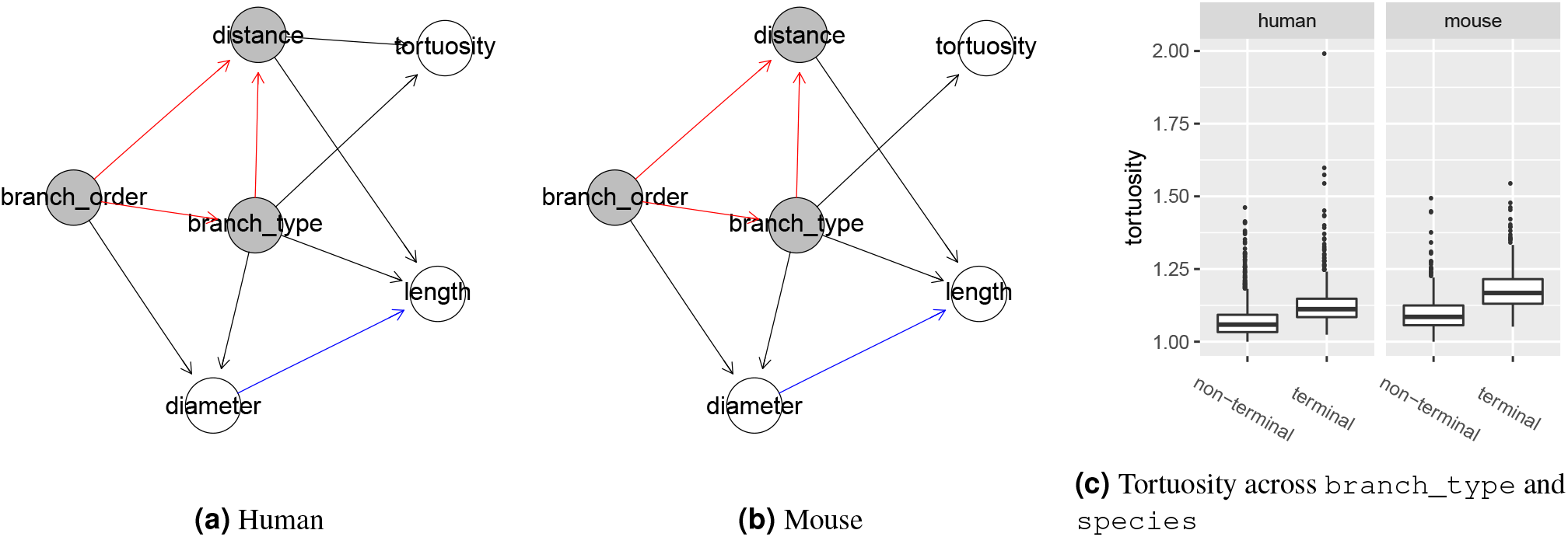
Bayesian networks learned from terminal and non-terminal branches of human data (a), mouse data (b), and differences in magnitude of `tortuosity` between terminal and non-terminal branches across species (c). Proximity between two nodes of a graph is unrelated to the strength of their correlation. The nodes of the morphological determinants are shaded in grey with red borders. non-terminal branches were in both human and mouse. Arcs among morphological determinants are depicted in red; among morphometrics in blue; the rest in black.

## 4 Discussion

A number of differences between terminal (Section 3.2)and non-terminal branches (Section 3.3) were consistent across species. First, there were less arcs per node in the Bayesian networks for the terminal branches, meaning that there were more independencies among variables in the terminal branches. For example, while longer non-terminal branches were thinner and more tortuous than the short ones, branch length was uncorrelated with diameter and tortuosity in terminal branches. Second, the conditional distributions of length and diameter were well approximated by a normal distribution for any branching order in the terminal branches yet for relatively few in in the non-terminal ones (see Table 2 in the Supplementary Information). These findings might suggest that there are fewer constraints on the growth of terminal branches, in both species, leading to little interaction among branch length, diameter and tortuosity.

Another shared difference between terminal and non-terminal branches was the effect of branch order and the distance from the soma on branch length. Indeed, the only morphometric marginally correlated with the distance from the soma and branch order in all four species-specific Bayesian networks (i.e., Figure 3a, Figure 3b, Figure 5a and Figure 5b) was branch length. The direction of the effect differed, however, as the branch length decreased with distance from the soma for terminal branches (Figure 3d) whereas it slightly increased for non-terminal branches (see Figure 2 in in the Supplementary Information).

Also, in terminal branches, the distance from soma was more informative about branch length than branch order, as length was independent of branch order given distance in both species (Figure 3a and Figure 3b), whereas in non-terminal branches distance was independent, conditional on branch order, of all morphometric in the mouse network (Figure 5b) and of 3 out of 5 morphometrics in the human network (Figure 5a). A possible explanation is that the wide range of distances within a branch order of terminal branches (up to 211.81 *μm* in Figure 4), relative to the narrow range of distances for non-terminal branches (up to 66.93 *μm* in Figure 6e), allows for variability in length within a branching order. Thus, when analysing terminal branches it is informative to consider the distance from the soma as complementary information to branch order whereas for non-terminal branches accounting for branch order alone may suffice.

The Bayesian networks of terminal branches were more similar between species than those of the non-terminal branches. Nonetheless, inter-species differences in the magnitude of length and tortuosity were more pronounced in terminal branches than in non-terminal ones, as evidenced by the lack of difference at some branch orders in the latter (Section 3.3). A similar result has been observed regarding the branch length of basal dendrites of temporal cortex pyramidal neurons of the human and the mouse13.

This paper introduced Bayesian networks as a multivariate model for comparison between species. Multivariate models are useful for inter-species comparison because we need to isolate heterogeneity that is due to intrinsic differences between species from the heterogeneity that is due to differences in morphological determinants such as branch order and branch type. By learning Bayesian networks from data we obtained concise graphical representations of the branch-level morphology in terms of probabilistic relationships among morphometrics and determinants. This representation allowed us to easily identify some differences and similarities between species. For example, the presence of an arc from branch order to distance suggested that the distance from soma might differ between the species within a branch order, which we then confirmed with Kruskal-Wallis tests, while a Kruskal-Wallis test that did not take branch order into account would have missed this difference. We found that determinants, such as branch order, were correlated with most morphometrics even after accounting for the species, and thus ought to be taken into account when testing for location differences. The determinants did not suffice, however, to fully explain any of the differences between species. For example, the lower tortuosity of human branches was not only due to their larger length. In addition to network structure, the signs and magnitudes of the regression coefficients in the conditional distributions of the networks indicated the direction and magnitude of the probabilistic relationships. For example, we found that the correlation of length and distance from soma had an opposite sign in terminal and non-terminal branches. The networks also uncovered interesting independencies and correlations among the variables. For example, the tortuosity was independent of branch length in terminal branches, although we might expect longer branches to be more tortuous, as in non-terminal branches.

Provided that our assumptions hold (see Section 2.6), the learned Bayesian networks are faithful models of the branch-level morphologies for the two species. As such, they could be used for more purposes than comparison. For example, to generate synthetic branches (see Ref.^25^) or to perform probabilistic queries about the morphology.

## 5 Data availability

The data analyzed in the current study are being published at a public repository. In the mean time, they are available from the corresponding author upon request.

## 6 Acknowledgements

This work has been partially supported by the Spanish Ministry of Economy and Competitiveness through the TIN2016-79684-P project. This project has received funding from the European Union’s Horizon 2020 Framework Programme for Research and Innovation under the Specific Grant Agreement No. 785907 (Human Brain Project SGA2).

## 7 Author contributions statement

B.M. designed and conducted the analysis and wrote the manuscript. R.B.P. and J.D.F. provided the data. All authors substantially reviewed the manuscript.

## 8 Additional information

The authors declare no competing interests.

