## Supplementary Information for "Comparing basal dendrite branches in human and mouse hippocampal CA1 pyramidal neurons with Bayesian networks"

### ABSTRACT

### 1 Morphometrics' conditional distributions

| Branch type | Species | Tortuosity | Diameter | Distance | Length | RBA | RTA |
|---|---|---|---|---|---|---|---|
| non-terminal | human | 0.00 | 0.00 | 0.00 | 0.00 | 0.16 | 0.03 |
| non-terminal | mouse | 0.00 | 0.00 | 0.00 | 0.00 | 0.53 | 0.08 |
| terminal | human | 0.00 | 0.08 | 0.00 | 0.04 | | |
| terminal | mouse | 0.00 | 0.32 | 0.03 | 0.92 | | |

**Table 1.** p-values for the Kolmogorov-Smirnov test of normality. Columns 1-2 indicate the groups and columns 3-8 show the p-values. RBA = remote bifurcation angle. RTA = remote tilt angle. Missing entries indicate that there were no observations for a given group.

| Branch type | Species | BO | Tortuosity | Diameter | Distance | Length | RBA | RTA |
|---|---|---|---|---|---|---|---|---|
| non-terminal | human | 0 | 0.00 | 0.00 | | 0.00 | 0.25 | 0.35 |
| non-terminal | human | 1 | 0.00 | 0.01 | 0.06 | 0.00 | 0.30 | 0.15 |
| non-terminal | human | 2 | 0.00 | 0.07 | 0.26 | 0.00 | 0.31 | 0.51 |
| non-terminal | human | 3 | 0.00 | 0.25 | 0.57 | 0.00 | 0.33 | 0.14 |
| non-terminal | human | 4 | 0.51 | 0.29 | 0.88 | 0.16 | 0.60 | 0.76 |
| non-terminal | mouse | 0 | 0.20 | 0.12 | | 0.01 | 0.77 | 0.77 |
| non-terminal | mouse | 1 | 0.48 | 0.02 | 0.09 | 0.00 | 0.60 | 0.29 |
| non-terminal | mouse | 2 | 0.05 | 0.19 | 0.01 | 0.00 | 0.59 | 0.82 |
| non-terminal | mouse | 3 | 0.04 | 0.14 | 0.36 | 0.04 | 0.94 | 0.75 |
| non-terminal | mouse | 4 | 0.29 | 0.13 | 0.45 | 0.40 | 0.66 | 0.99 |
| terminal | human | 1 | 0.08 | 0.36 | 0.25 | 0.56 | | |
| terminal | human | 2 | 0.00 | 0.28 | 0.00 | 0.31 | | |
| terminal | human | 3 | 0.00 | 0.30 | 0.00 | 0.40 | | |
| terminal | human | 4 | 0.00 | 0.83 | 0.00 | 0.58 | | |
| terminal | mouse | 1 | 0.09 | 0.76 | 0.52 | 0.42 | | |
| terminal | mouse | 2 | 0.71 | 0.20 | 0.08 | 0.85 | | |
| terminal | mouse | 3 | 0.11 | 0.64 | 0.14 | 0.86 | | |
| terminal | mouse | 4 | 0.04 | 0.36 | 0.79 | 0.95 | | |

**Table 2.** p-values for the Kolmogorov-Smirnov test of normality. Columns 1-3 indicate the groups and columns 4-9 show the p-values. RBA = remote bifurcation angle. RTA = remote tilt angle. Missing entries indicate that there were no observations for a given group.

### 2 Non-terminal branches: correlation of distance from soma and morphometrics

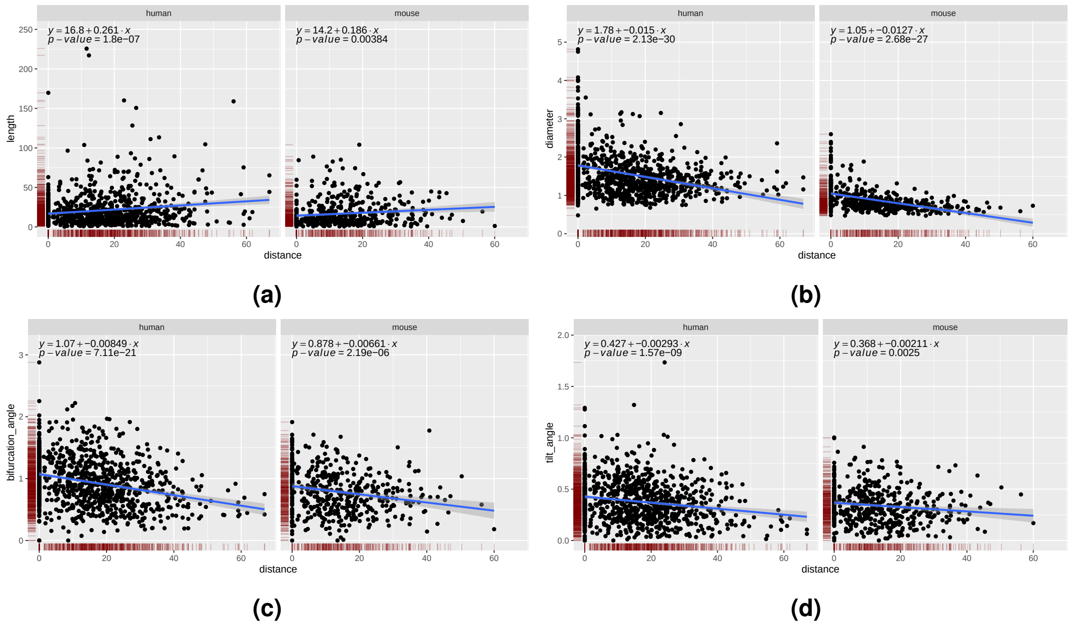


**Figure 1.** Non-terminal branches. Linear correlation of `distance` and `length`, `diameter`, `bifurcation_angle`, and `tilt_angle`, respectively. The sign of the correlation coefficient was the same for both species in all cases, with the regression lines roughly parallel.

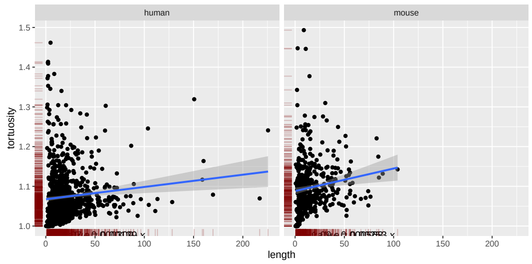


**(a)**

**Figure 2.** Non-terminal branches. `tortuosity` as a function of `length`.

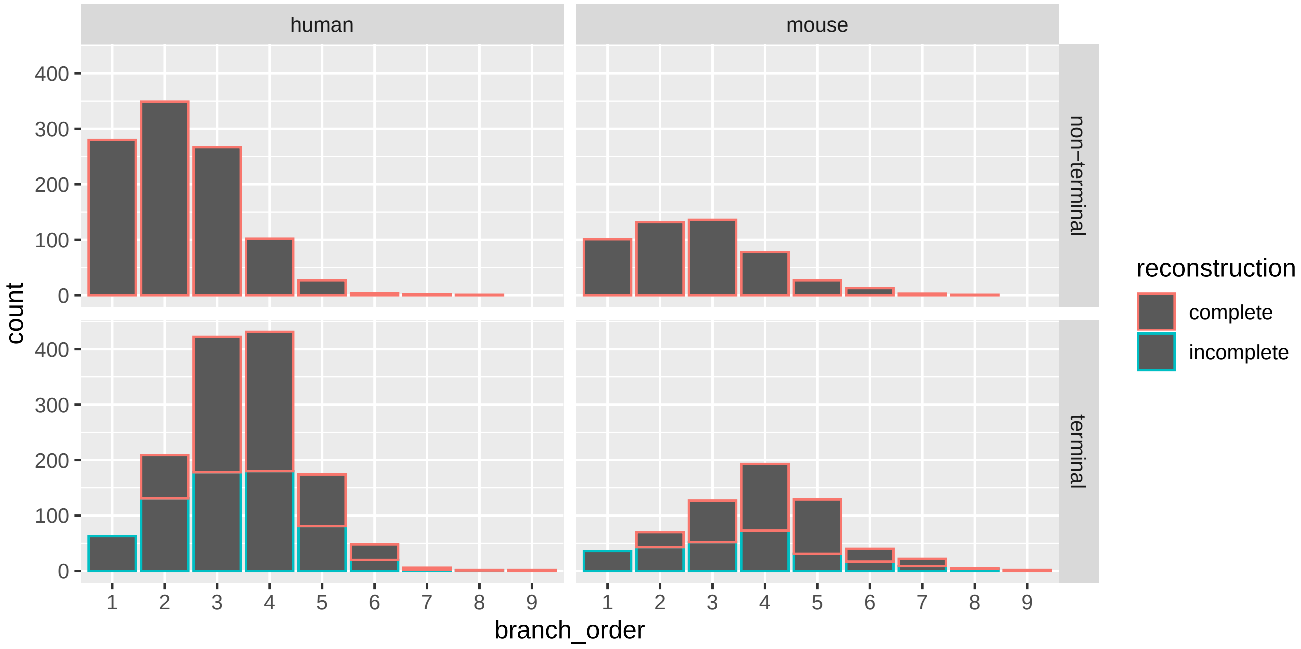


**Figure 3.** Complete and incomplete branch frequencies per species, branch type, and branching order.

### 3 Complete and incomplete branches

### 4 Non-terminal branches: inter-species differences when conditioning on branch order

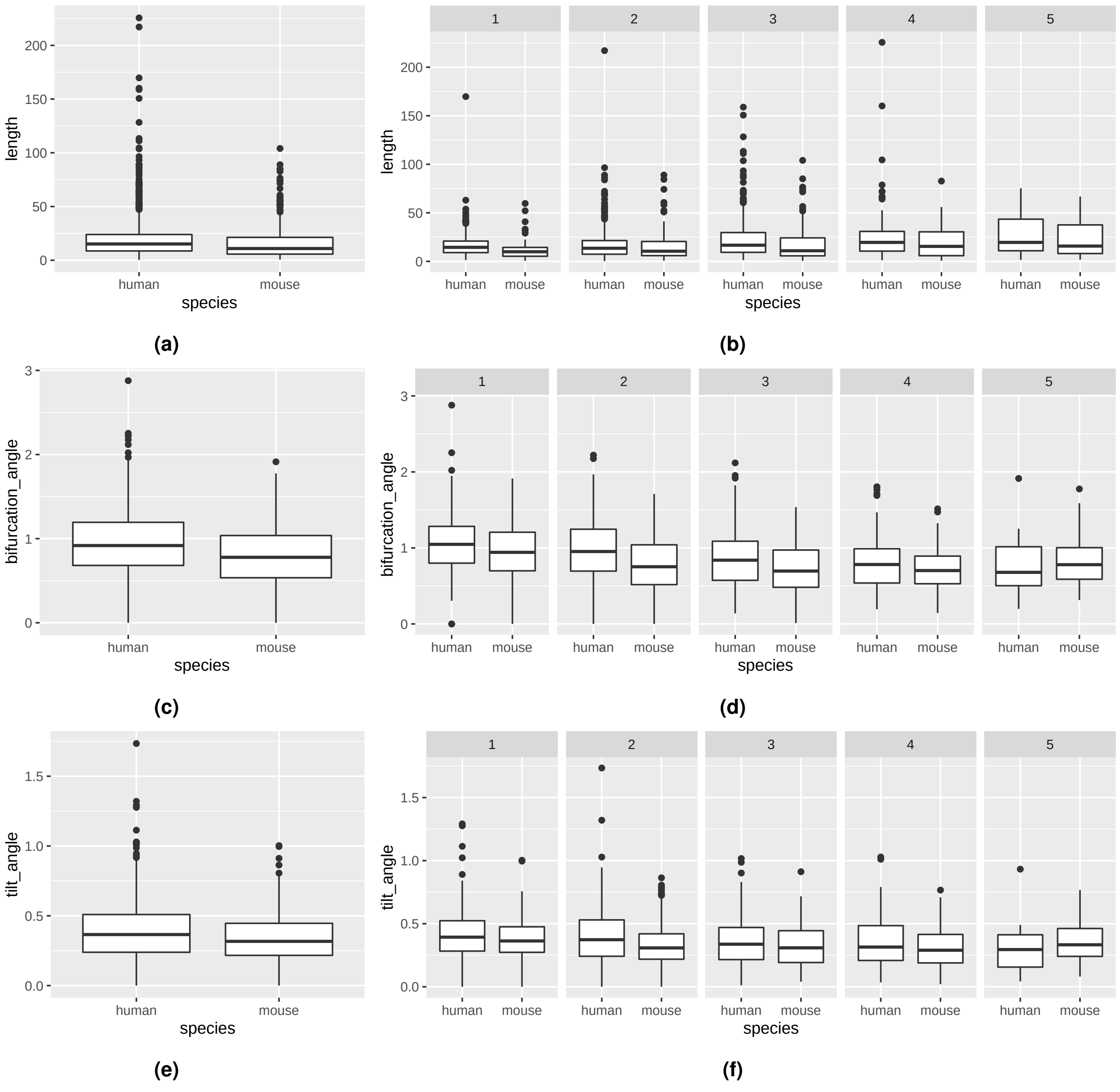


**Figure 4.** Differences in magnitude of variables of non-terminal branches between the species. (a) `length`: Marginal KW p-value 1.793e-07. KW p-values 6.71e-07, 0.08872, 0.0004447, 0.05116, and 0.4398 for branch orders 1, 2, 3, 4 and 5, respectively; (b) `bifurcation_angle`: Marginal KW p-value 6.841e-13. KW p-values 0.02463, 1.653e-06, 0.0004749, 0.142, and 0.5848 for branch orders 1, 2, 3, 4 and 5, respectively; (c) `tilt_angle`: Marginal KW p-value 0.000121. KW p-values 0.1185, 0.003194, 0.3144, 0.1535, and 0.3001 for branch orders 1, 2, 3, 4 and 5, respectively.
